# Using machine learning and big data to explore the drug resistance landscape in HIV

**DOI:** 10.1101/2021.03.16.435621

**Authors:** Luc Blassel, Anna Tostevin, Christian Julian Villabona-Arenas, Martine Peeters, Stéphane Hué, Olivier Gascuel, On behalf of the UK HIV Drug Resistance Database

## Abstract

Drug resistance mutations (DRMs) appear in HIV under treatment pressure. DRMs are commonly transmitted to naive patients. The standard approach to reveal new DRMs is to test for significant frequency differences of mutations between treated and naive patients. However, we then consider each mutation individually and cannot hope to study interactions between several mutations. Here, we aim to leverage the ever-growing quantity of high-quality sequence data and machine learning methods to study such interactions (i.e. epistasis), as well as try to find new DRMs.

We trained classifiers to discriminate between Reverse Transcriptase Inhibitor (RTI)-experienced and RTI-naive samples on a large HIV-1 reverse transcriptase (RT) sequence dataset from the UK (*n ≈* 55, 000), using all observed mutations as binary representation features. To assess the robustness of our findings, our classifiers were evaluated on independent data sets, both from the UK and Africa. Important representation features for each classifier were then extracted as potential DRMs. To find novel DRMs, we repeated this process by removing either features or samples associated to known DRMs.

When keeping all known resistance signal, we detected sufficiently prevalent known DRMs, thus validating the approach. When removing features corresponding to known DRMs, our classifiers retained some prediction accuracy, and six new mutations significantly associated with resistance were identified. These six mutations have a low genetic barrier, are correlated to known DRMs, and are spatially close to either the RT active site or the regulatory binding pocket. When removing both known DRM features and sequences containing at least one known DRM, our classifiers lose all prediction accuracy. These results likely indicate that all mutations directly conferring resistance have been found, and that our newly discovered DRMs are accessory or compensatory mutations. Moreover, we did not find any significant signal of epistasis, beyond the standard resistance scheme associating major DRMs to auxiliary mutations.

**Author summary:** Almost all drugs to treat HIV target the Reverse Transcriptase (RT) and Drug resistance mutations (DRMs) appear in HIV under treatment pressure. Resistant strains can be transmitted and limit treatment options at the population level. Classically, multiple statistical testing is used to find DRMs, by comparing virus sequences of treated and naive populations. However, with this method, each mutation is considered individually and we cannot hope to reveal any interaction (epistasis) between them. Here, we used machine learning to discover new DRMs and study potential epistasis effects. We applied this approach to a very large UK dataset comprising *≈* 55, 000 RT sequences. Results robustness was checked on different UK and African datasets.

Six new mutations associated to resistance were found. All six have a low genetic barrier and show high correlations with known DRMs. Moreover, all these mutations are close to either the active site or the regulatory binding pocket of RT. Thus, they are good candidates for further wet experiments to establish their role in drug resistance. Importantly, our results indicate that epistasis seems to be limited to the classical scheme where primary DRMs confer resistance and associated mutations modulate the strength of the resistance and/or compensate for the fitness cost induced by DRMs.

## Introduction

Drug resistance mutations (DRMs) arise in Human Immunodeficiency Virus-1 (HIV-1) due to antiretroviral treatment pressure, leading to viral rebound and treatment failure [1, 2]. Furthermore, drug-resistant HIV strains can be transmitted to treatment-naive individuals and further spread throughout the population over time [3–5]. These transmitted resistant variants limit baseline treatment options and have clinical and public health implications worldwide. Almost all drugs to treat HIV target the reverse transcriptase (RT), encoded by the *pol* gene. Lists of DRMs are regularly compiled and updated by experts in the field, based on genotype analyses and phenotypic resistance tests or clinical outome in patients on ART [6–8]. However, with the developement of new antiretroviral drugs that target RT but also other regions of the *pol* gene like protease or integrase, and the use of anti-retrovirals in high risk populations by pre-exposure prophylaxis (PREP), it is important to further our understanding of HIV polymorphisms and notably the interactions between mutations and epistatic effects.

Among known DRMs, some mutations, such as M184V, directly confer resistance to antiretrovirals, more precisely the commonly used NRTI, 3TC (lamivudine) and FTC (emtricitabine) and are called primary or major drug resistance mutations, while some mutations like E40F have an accessory role and increases drug resistance when appearing alongside primary DRMs. Moreover, some mutations like S68G seem to have a compensatory role, but are not known to confer any resistance nor modulate resistance induced by primary DRMs. All of these mutations might have different functions in the virus, but they are all known to be associated with drug resistance phenomena. Therefore, during the rest of this article we will refer to all of these known mutations as resistance associated mutations (RAMs), rather than DRMs which is too specific, and our goal will be to search for new RAMs and study the interactions between known RAMs and the new ones.

Classically, new RAMs have been found using statistical testing and large multiple sequence alignments (MSA) of the RT [9, 10]. Tests are performed for mutations of interest on a given MSA to check if they are associated with the treatment status and outcome of the individual the viral sequences were sampled from. The test significance is corrected for multiple testing as all mutations associated to every MSA position is virtually a resistance mutation and tested. After this preliminary statistical search, the selected mutations are scrutinized to remove the effects of phylogenetic correlation (i.e. typically counting two sequences which are identical or closely related due to transmission rather than independent acquisition twice [11]) and check that the same mutation occurred several times in different subtypes and populations being treated with the same drug. Then, these mutations can be further experimentally tested in vitro or in vivo to validate phenotypic resistance. This method has worked well, but by design it is incapable of studying the effect of several mutations at once, since if we have to test all couples or triplets of mutations, we quickly lose all statistical power when correcting for multiple testing [12], due to the sheer number of tests to perform.

The goal of this study is two-fold. On the one hand we aim to find new potential RAMs by applying multiple testing and a machine learning based approach to a very large HIV-1 RT sequence datasets from the UK (*n ≈* 55, 000) (http://www.hivrdb.org.uk/) and Africa (*n ≈* 4, 000) [10]. On the other hand, we aim to detect associations between mutations and their effect on antiretroviral resistance in order to study potential underlying epistasis. The African and UK datasets are very different both on genetic and treatment history standpoints, therefore training classifiers on the UK dataset and testing them on the African one, should guaranty the robustness of our findings and greatly alleviate phylogenetic correlation effects. In the following sections, we first describe the data then the methods used. Our results include the assessment of the performance of our classifiers even when trained on data devoid of any known resistance-associated signal; as well as a description of the main features (prevalence and correlation to known mutations, genetic barrier and structural analysis) of six potentially resistance associated mutations, newly discovered thanks to our approach. These results and perspectives are discussed in the concluding section.

## Materials and methods

### Data

In this study, we used all the mutations that appeared in the Stanford HIV Drug resistance database, both for NRTI (Nucleoside Reverse Transcriptase Inhibitors https://hivdb.stanford.edu/dr-summary/comments/NRTI/) and NNRTI (Non Nucleoside RTI https://hivdb.stanford.edu/dr-summary/comments/NNRTI/) as known RAMs. To discover new RAMs, assess their statistical significance and study potential epistatic effects, we used two datasets of HIV-1 RT sequences. A large one (*n* = 55, 539) from the UK HIV Drug Resistance Database (http://www.hivrdb.org.uk/) and a smaller (*n* = 3, 990) one from 10 different western, eastern and central African countries [10]. In the UK dataset, sequences from RTI-naive individuals formed the majority class with 41,921 sequences (75%). In the African dataset, both classes were more balanced with 2,316 RTI-naive sequences (58%). In the UK dataset, RTI-naive sequences had at least one known RAM in 25% of cases most likely due to transmissions to naive patients or undisclosed treatment history, against 48% in RTI-experienced sequences, thus making the discrimination between the RTI-experienced and RTI-naive sequences particularly difficult. In the African dataset this distribution was more contrasted, with only 14% of RTI-naive sequences having at least one known RAM, versus 83% of RTI-experienced sequences. The African dataset was also much more genetically diverse with 24 different subtypes and CRFs compared to the 2 subtypes (B and C) that we retained for this study from the UK cohort. The majority of the sequences from the African dataset were samples from Cameroon (27%), Democratic Republic of Congo (17%), Burundi (15%), Burkina Faso (13%) and Togo (11%).

It is important to note that RTI-experienced sequences in both of these datasets can be considered as resistant to treatment. Since the viral load was sufficiently high to allow for sequencing of the virus, we can consider that the ART has failed. However, in some cases this resistance is caused by non adherence to ART, rather than by the presence of RAMs, therefore adding some noise to the relationship between treatment status and resistance.

In addition to differences in size, balance between RTI-naive and experienced classes, and the genetic difference between the UK and African datasets, there are also significant differences resulting from differing treatment strategies. In the UK and other higher income countries, the treatment is often tailored to the individual with genotype testing, which result in specific treatment as well as thorough follow-ups and high treatment adherence. In the African countries of the dataset that we used, the treatment is ZDV/ d4T (NRTI) + 3TC (NRTI) + NVP/EFV (NNRTI) in most cases [10], and this treatment is generalized to the affected population, with poorer follow-up and adherence than in the UK. This discrepancy could lead to different mutations arising in both datasets, however since the treatment strategy is a combination of both NRTI and NNRTI drug classes, as in many countries, similar RAMs arise [10]. Furthermore, there is potentially more uncertainty in the African dataset than in the UK. For example some individuals may have unofficially taken antiretroviral drugs but still identify themselves as RTI-naive or report having some form of ART while not having been treated for HIV [13]. All of this explains the high prevalence of multiple resistance in the African data set: the median number of RAMs in sequences containing at least one RAM is 3 in the African sequences, while it is 1 in UK sequences (Table 1). Thus, we can say that African sequences are highly resistant, with possibly different mutations and epistatic effects, compared to their UK counterparts.

**Table 1.** Summary of the UK and African datasets.

|  |  | UK |  | Africa |  |
| --- | --- | --- | --- | --- | --- |
| size |  | 55539 |  | 3990 |  |
| RTI naive | with known RAMs | 11429 | (21%) | 318 | (8%) |
|  | without known RAMs | 30492 | (55%) | 1998 | (50%) |
| RTI experienced | with known RAMs | 6633 | (12%) | 1388 | (35%) |
|  | without known RAMs | 6985 | (13%) | 286 | (7%) |
| sequences with $\geq 2$ known RAMs | | 8034 | (14%) | 1308 | (33%) |
| max known RAM number |  | 13 |  | 17 |  |
| Median known RAM number |  | 1 |  | 3 |  |
| number of subtypes / CRFs |  | 2 |  | 24 |  |
| subtypes / CRFs | A | 0 | (0%) | 472 | (12%) |
|  | B | 37806 | (68%) | 64 | (2%) |
|  | C | 17733 | (32%) | 702 | (18%) |
|  | CRF02_AG | 0 | (0%) | 1477 | (37%) |

Percentages are computed with regards to the size of the considered dataset (e.g. 21% of the sequences of the UK dataset are RTI-naive and have at least one known RAM). The median number of RAMs was computed only on sequences that had at least one known RAM.

All these differences between the two datasets helped us to assess the generalizability of our method and the robustness of the results. That is to say, if signal extracted from the UK dataset was still relevant on such a different dataset as the African one, we could be fairly reassured in regard to the biological and epidemiological relevance of the observed signal.

Sequences in both African and UK datasets were already aligned. We used the Sierra web service (https://hivdb.stanford.edu/page/webservice/) to get amino acid positions relative to the reference HXB2 HIV genome. This allowed us to determine all the amino acids present at each reference position in both datasets, among which we distinguished the “reference amino acids” for each position, corresponding to the B and C subtype reference sequences obtained from the Los Alamos sequence database (http://www.hiv.lanl.gov/). All the other, non-reference amino acids are named “mutations” in the following, and the set of mutations was explored to reveal new potential RAMs.

To train our supervised classification methods [14–16], the sequence data needed to be encoded to numerical vectors. We represented each sequence by a binary vector denoting the presence/absence of every mutation present in the UK dataset. Therefore, each binary representation feature corresponded to a specific mutation seen in the dataset. Some of these features corresponded to known RAMs and are named RAM features in the following.

### Classifier training

In order to find new potential RAMs, we first followed the conventional multiple testing approach [10]. We first used Fisher exact tests to identify which of these mutations were significantly associated with anti-retroviral treatment. All the resulting p-values were then corrected for multiple testing using the Bonferroni correction [17]. Those for which the corrected p-value was *≤* 0.05 were then considered as significantly associated with treatment and potentially implicated in resistance.

This method was complemented by our parallel, machine learning based approach. In order to extract potential RAMs, we trained several classifiers to discriminate between RTI-experienced and RTI-naive sequences represented by the binary vectors described above. Once the classifiers were trained, we extracted the most important representation features, which corresponded to potentially resistance-associated mutations (PRAM in short). To this aim we chose three interpretable supervised learning classification methods so as to be able to extract those features:

1. Multinomial naive Bayes (NB), which estimates conditional probabilities of being in the RTI-experienced class given a set of representation features [18]; the higher (*≈* 1.0) and the lower (*≈* 0) conditional probabilities correspond to the most important features.
2. Logistic regression (LR) with L1 regularization (LASSO) [14] which assigns weights to each of the features, whose sign denotes the importance to one of the 2 classes, and whose absolute value denotes the weight of this importance.
3. Random Forest (RF), which has feature importance measures based on the Gini impurity in the decision trees [19].

Interpretability was the main driver behind our classification method choice, and the reason we chose not to use more over-complicated, “black-box” methods which are hard to interpret [20–22]. Moreover, these three classification methods have the potential to detect epistatic effects. With RF, the discrimination is based on the combination of a few features (i.e. mutations), while with LR the features are weighted positively or negatively, thus making it possible to detect cumulative effects resulting from a large number of mutations, which individually have no discrimination power. Naive Bayes is a very simple approach, generally fairly accurate, and in between the two others in terms of explanatory power [16]

In order to be able to compare all these approaches in a common framework, we devised a very simple classifier out of the results of the Fisher exact tests. This “Fisher” classifier (FC) predicts a sequence as RTI-experienced if it has at least one of the mutations significantly associated to treatment. In this way we were able to compute metrics for all classification methods and compare their performance.

It is important to note that in all of these approaches we chose to discriminate RTI-naive from RTI-experienced sequences, regardless of the type of RTI received. A first reason is that we did not have detailed enough treatment history for sequences in the UK and African datasets. Moreover, even without segmenting by treatment type, the size of the training set and the power of our classification methods were both high enough to be able to detect all kinds of resistance associated mutations. We shall see (Result section) that we were able to determine the likely treatment involved by further examining the important extracted features and comparing them to known RAMs. Furthermore, since the treatment strategies are so different between the UK and African sequences, training on sequences having received different treatments should increase the robustness of our classifiers and the relevance of the mutations selected as potentially associated to resistance.

To avoid phylogenetic confounding factors (e.g. transmitted mutations within a specific country of region), and avoid finding mutations potentially specific to a given subtype, we split the training and testing sets by HIV-1 M subtype. This resulted in training a set of classifiers on all subtype B sequences of the UK dataset and testing them on subtype C sequences from the UK dataset, training another set of classifiers on the subtype C sequences of the UK dataset and testing on the subtype B sequences from the UK dataset, as well as training a final set of classifiers on the whole UK dataset, but testing it on the smaller African dataset with a completely different phylogenetic makeup and treatment context [10]. Furthermore, in order to identify novel RAMs and study the behavior of the classifiers, we repeated this training scheme on both datasets, each time removing resistance-associated signal incrementally: first by removing all representation features corresponding to known RAMs from the dataset, and second by removing all sequences that had at least one known RAM. This resulted in each type of classifier being trained and tested 9 times, on radically different sets to ensure the interpretability and robustness of the results (see Table 2).

**Table 2.** All training and testing datasets used during this study.

| Signal removal level | Trained on |  | Tested on |  |
| --- | --- | --- | --- | --- |
| None | UK, subtype B | (37806) | UK, subtype C | (17733) |
|  | UK, subtype C | (17733) | UK, subtype B | (37806) |
|  | UK, subtypes B & C | (55539) | Africa, all subtypes | (3990) |
| Known RAM features removed | UK, subtype B | (37806) | UK, subtype C | (17733) |
|  | UK, subtype C | (17733) | UK, subtype B | (37806) |
|  | UK, subtypes B & C | (55539) | Africa, all subtypes | (3990) |
| Known RAM features & sequences with $\geq 1$ known RAM removed | UK, subtype B | (24422) | UK, subtype C | (24422) |
|  | UK, subtype C | (13055) | UK, subtype B | (13055) |
|  | UK, subtypes B & C | (37477) | Africa, all subtypes | (2284) |

The number of sequences in each dataset is shown in parentheses

### Measuring classifier performance

To compare the performance of our classifiers we used balanced accuracy [23], which is the average of accuracies (i.e. percentages of well-classified sequences) computed separately on each class of the test set. This score takes into account, and corrects for, the imbalance between RTI-naive and RTI-experienced samples, which would lead to a classifier always predicting a sequence as RTI-naive getting a classical accuracy score of up to 77% (i.e. the frequency of naive sequences in the UK dataset). We also computed the adjusted mutual information (AMI) between predicted and true sequence labels, which is a normalized version of MI allowing comparison of performance on differently sized test sets [24]. Additionally, mutual information (MI) was used to compute p-values and assess the significance of the classifiers’ predictive power. The probabilistic performance of the classifiers was evaluated using an adapted Brier score [15] more suited to binary classification, which is the mean squared difference between the actual class (coded by 1 and 0 for the RTI-experienced and RTI-naive samples respectively) and the predicted probability of being RTI-experienced. This approach refines the standard accuracy measure by rewarding methods that well approximate the true status of the sample (eg. predicting a probability of 0.9 while the true status is 1); conversly, binary methods (predicting 0 or 1, but no probabilities) will be penalized if they are often wrong. The Brier approach thus assigns better scores to methods that recognize their ignorance than to methods producing random predictions.

## Results

### Classifier performance & interpretation

As can be seen in Fig 1a and b, when all RAM features and sequences were kept in the training and testing sets, classifiers had good prediction accuracy, with the machine learning classifiers slightly outperforming the “Fisher” classifier. When removing RAM features from the training and testing sets, the classifiers retained a significant prediction accuracy, especially with the African data set and its multiple RAMs that are observed in a large number of sequences (but removed in this experiment). In this configuration the ML classifiers had a similar performance to the “Fisher” classifier, except for the random forest that is slightly less accurate, likely due to overfitting. Also, when removing sequences that had known RAMs, every classifier lost all prediction accuracy, and none could distinguish RTI-naive from RTI-experienced sequences. Regarding the Brier sore, we see the advantage of the machine learning classifiers over the “Fisher” classifier, which is worse than random predictions when known RAMs are removed. The ability of machine learning classifiers to quantify the resistance status should be an asset for many applications.

**Fig 1.**
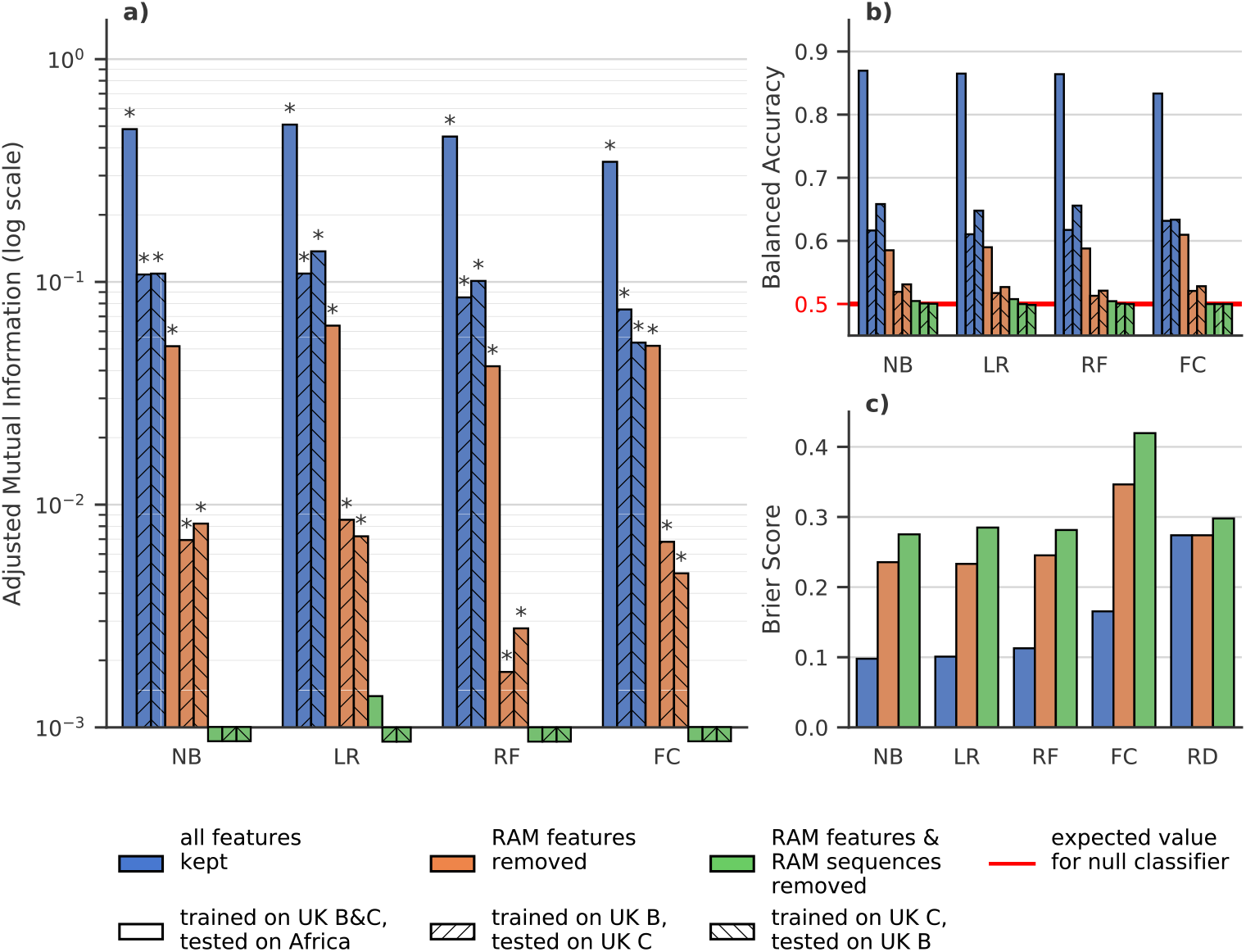
Classifier Performance on UK and African datasets. **NB**: naive Bayes, **LR**: Logistic Regression with Lasso regularization, **RF**: Random Forest, **FC**: Fisher Classifier, **RD**: Agnostic random probabilistic classifier (this classifier predicts, as the probability of a sample belonging to a class, the frequency of that class in the training data). **a)** Adjusted mutual information (higher is better) between ground truth and predictions by classifiers trained on dataset with all features (blue), without features corresponding to known RAMs (orange) and without RAM features and without sequences that have at least 1 known RAM (green). Hatching indicates the training set on which a classifier was trained and the testing set on which the performance was measured. The expected value for a null classifier is 0, and 1 for a perfect classifier and a * denotes that the p-value derived from mutual information is ≤ 0.05. For example when trained with all features all the classifiers have a significative MI. Conversly when removing RAM features and RAM sequences none of the classifiers have a significative MI and only LR trained on the entirety of the UK dataset has an AMI *>* 10^*-*3^ **b)** Balanced Accuracy score, i.e. average of accuracies per-class (higher is better) for the same classifiers as in a). The red line at *y* = 0.5 is the expected balanced accuracy for a null classifier that only predicts the majority class as well as a random uniform (i.e. 50/50) classifier. **c)** Brier score, which is the mean squared difference between the sample’s experience to RTI and the predicted probability of being RTI experienced.(lower is better), for the same classifiers as in **a)** and **b)**.

The fact that classifiers retained prediction accuracy after removing known RAM corresponding features suggests that there was some residual, unknown resistance-associated signal in the data. The fact that this same power was non-existent when removing the known RAM-containing sequences from the training and testing sets indicates that this residual signal was contained in these already mutated sequences. This suggests that the mutations that are found in the RAM removed experiment (see list below) are most likely accessory mutations that accompany known RAMs. This also suggests that all primary DRMs, i.e. that directly confer antiretroviral resistance, have been identified, which is reassuring from a public health perspective.

The performance discrepancy between the UK and African test sets can be explained by several factors. Firstly, African sequences that have known RAMs are more likely to have multiple RAMs, and thus more (known and unknown) resistance-associated features than their UK counterparts (c.f. Table 1). This means that resistant African sequences are easier to detect even when removing known RAMs. Secondly, RTI-naive sequences in the UK test sets are more likely to have known RAMs than their African counterparts (c.f. Table 1) and therefore more companion mutations. This means that the RTI-naive sequences in the UK test set are more likely to be misclassified as RTI-experienced than in the African test set.

### Additional classification results

The fact that, when looking at classifiers trained without known RAMs, “Fisher” classifiers perform as well as the machine learning ones, leads us to believe that there is little interaction between mutations that would explain resistance better than taking each mutation separately. It is therefore likely that the kind of epistatic phenomena we were looking for, combining several mutations that do not induce any resistance when taken separately, do not come into play here. We are in a classical scheme where primary DRMs confer resistance and associated mutations reinforce the strength of the resistance and/or compensate for the fitness cost induced by primary DRMs.

It is important to remember that in the previous section we were trying (as usual, e.g. see [10]) to find novel mutations associated with resistance by discriminating RTI-naive from RTI-experienced sequences, both with the statistical tests and the classifiers. However, this is intrinsically biased and noisy. Indeed, a RTI-naive sequence is not necessarily susceptible to RTIs as a resistant strain could have been transmitted to the individual. Conversely, an RTI-experienced sequence may not be resistant to treatment, due to poor ART adherence for example. We must therefore keep in mind that the noisy nature of the relationship between resistance and treatment status is partly responsible for the lower performance of classifiers trained on the UK sequences with reduced signal.

Moreover, as all the additional resistance signal we detected is associated to the sequences having at least one known RAM (see above), we performed another analysis trying to discriminate between the sequences having at least one known RAM and those having none. The goal was to check that the mutations we discovered by discriminating RTI-experienced from RTI-naive samples, are truly accessory and compensatory mutations. As can be seen in Fig 2 a and b, the classifiers trained to discriminate sequences that have at least one known RAM from those that have none, on datasets from which all features corresponding to known RAMs were removed, perform much better than classifiers trained to discriminate RTI-experienced from RTI-naive sequences. This increase in performance is especially visible for classifiers tested on UK sequences (more difficult to classify than the African ones, see above), with an AMI often almost one order of magnitude higher for the known-RAM presence/absence classification task. This further reinforces our belief that all there is a fairly strong residual resistance-signal in sequences that contain known RAMs, due to new accessory and compensatory mutations identified by our classifiers and Fisher tests. As a side note, Logistic regression (LR) consistently outperforms other classifiers, a tendency already observed in Fig 1.

**Fig 2.**
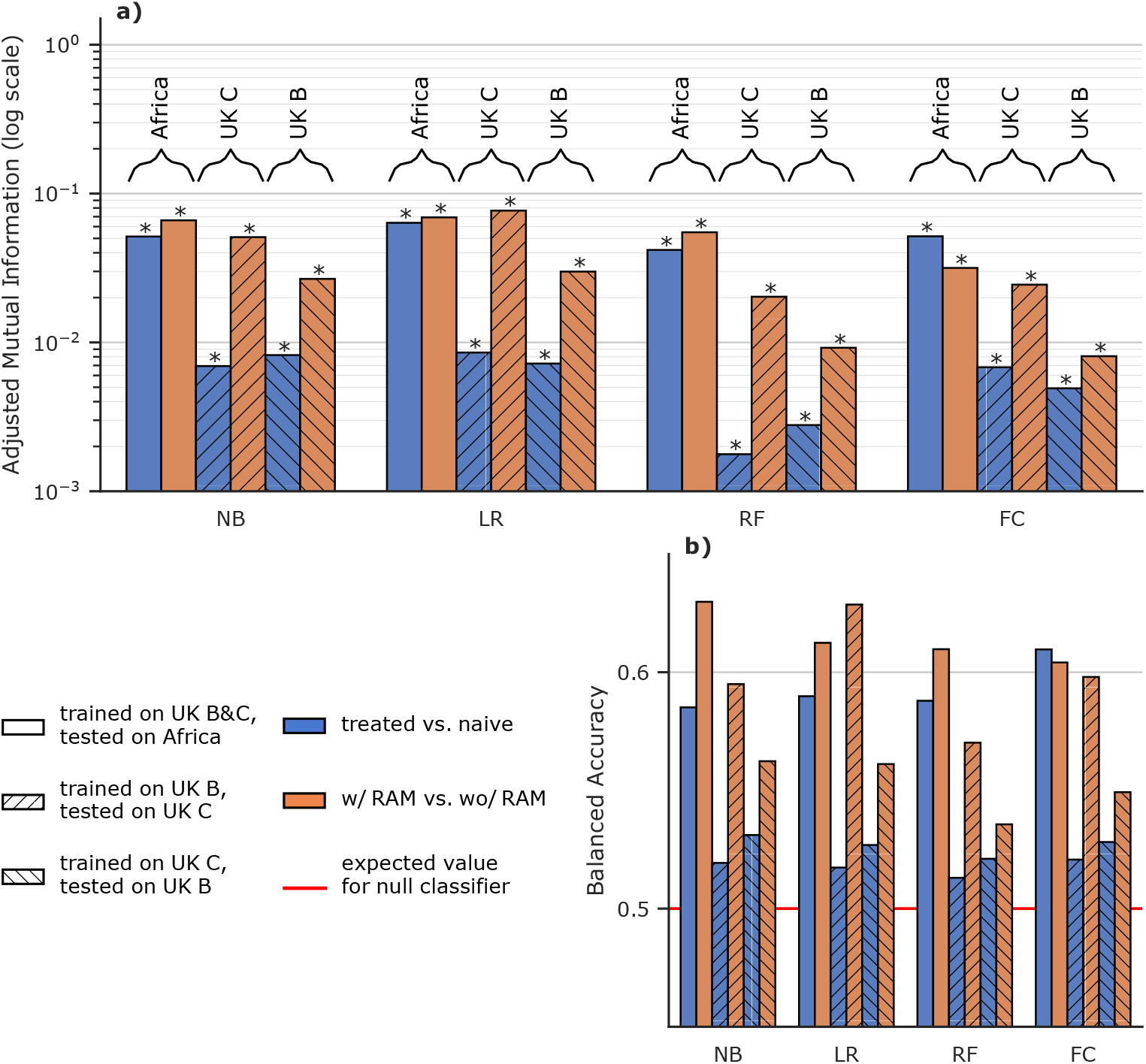
Discrimination between sequences having at least one RAM, and those having none on sequences with training features corresponding to known RAMs removed. **NB**: naive Bayes, **LR**: Logistic Regression with Lasso regularization, **RF**: Random Forest, **FC**: Fisher Classifier. **a)** Adjusted mutual information (higher is better) for classifiers trained without features corresponding to known RAMs. The classifiers are either trained to discriminate RTI-naive from RTI-experienced sequences (blue), or sequences with at least one known RAM from sequences that have none (orange). Hatching and braced annotations indicate the training and testing sets resulting in a given performance measure. **b)** Balanced accuracy, i.e. average of accuracies per-class for the same classifiers as in **a)** (higher is better). The red line at *y* = 0.5 is the expected value for a classifier only predicting the majority class as well as a random uniform (50/50) classifier.

### Identifying new mutations from classifiers

We assessed the importance of each mutation in the learned internal model of all the classifiers, in the setting where all known RAMs have been removed from the training dataset. For the Fisher classifier, we used one minus the p-value of the exact Fisher test as the importance value, therefore the more significantly associated mutations have the higher importance value and were ranked first. For a given classification task, we ranked each mutation according to the appropriate importance value for each classifier (see above), trained on the B or C subtypes, with the highest importance value having a rank of 0. We then computed the average rank for each mutation and each classification task (RTI-naive/RTI-experienced and RAM present/RAM absent). This gave us, for each classification task, a ranking of mutations potentially associated with resistance that took into account the importance given to this new mutation by each classifier trained on this task. Mutations that were in the 10 most important mutations for both of the classification tasks were considered of interest. Based on these criteria we selected the following potentially resistance-associated mutations (w.r.t. the HXB2 reference genome): L228R, L228H, E203K, D218E, I135L and H208Y. These mutations are referred to as “new mutations” in the rest of this study.

To check the epistatic nature of these selected mutations we defined the following ratio *ρ*(*new, X*) between a new mutation and a binary character *X*:

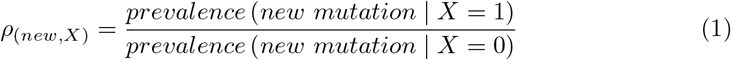

This ratio gives us a measure for how over-represented each of our new mutations is in sequences that have the *X* character compared to those that don’t.

To get a general idea of this over-representation, for each new mutation we computed *ρ*(*new, treatment*) comparing the prevalence of the new mutation in RTI-experienced and RTI-naive sequences. We also computed *ρ*(*new, withRAM*) comparing the prevalence the new mutation in sequences having at least one known RAM and sequences that have none. Both of these ratios are shown in Table 3 for each new mutation.

**Table 3.**
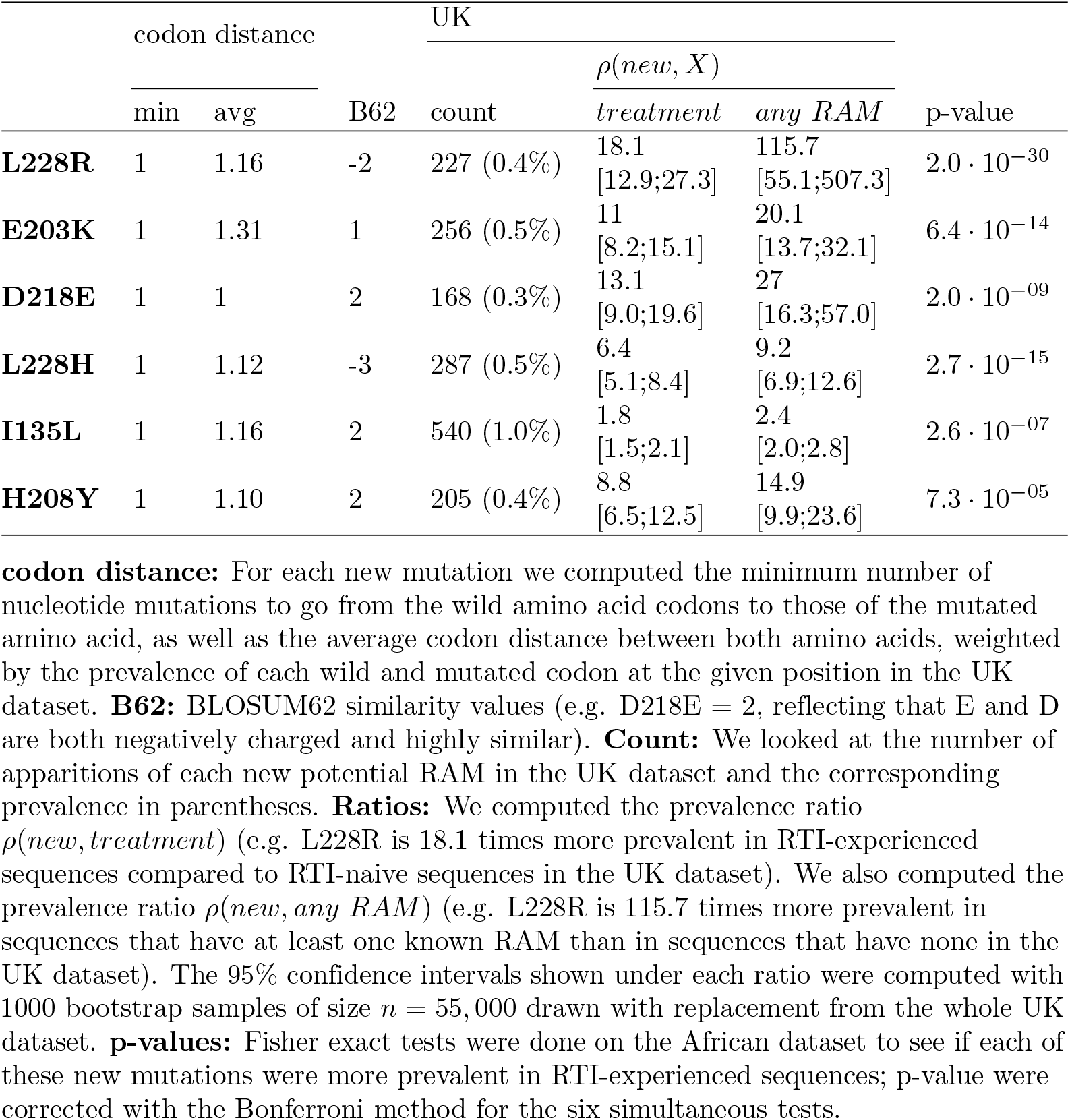
Analysis of new potential RAMs.

We then computed the *ρ*(*new, RAM*) ratios for each known RAM present in more than 0.1% of UK sequences and the new mutations. In Fig. 3 we see these ratios for which the lower bound of the 95% confidence interval, computed on 1000 bootstrap samples from the UK dataset, were greater than 4.

**Fig 3.**
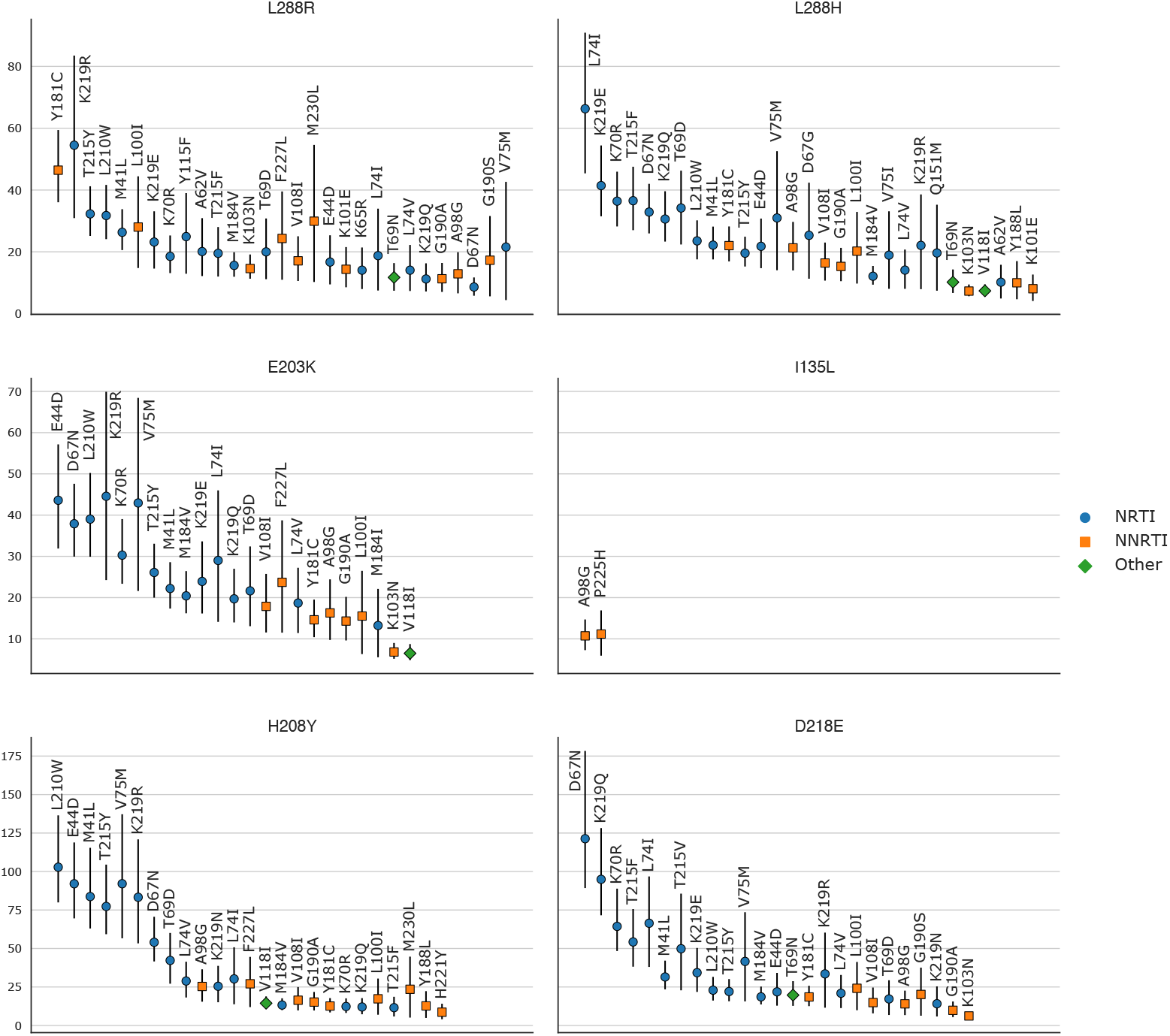
Prevalence ratios of the new mutations with regards to known RAMs on the UK dataset. (i.e. the prevalence of the new mutation in sequences with a given known RAM divided by the prevalence of the new mutation in sequences without this RAM). Ratios were only computed for mutations (new and RAMs) that appeared in at least 0.1% (=55) sequences. 95% confidence intervals, represented by vertical bars, were computed with 1000 bootstrap samples of UK sequences. Only ratios with a lower CI boundary greater than 4 are shown. The shape and color of the point represents the type of RAM as defined by Stanford’s HIVDB. Blue circle: NRTI, orange square: NNRTI, green diamond: Other. Ratio values are shown from left to right, by order of decreasing values on the lower bound of the 95% CI.

### Detailed analysis of potentially resistance-associated mutations

As can be seen in Table 3, all of these new mutations except for I135L, are highly over-represented in RTI-experienced sequences and sequences that already have known RAMs, with lower bounds on the 95% ratio CI always greater than 5, and often exceeding 10. When looking at the ratios computed for individual RAMs on the UK dataset (Fig 3), this impression is confirmed with very high over-representation of these new mutations potentially associated with resistance in sequences that have a given known RAM, with 95% ratio lower CI bounds sometimes greater than 80 (H208Y/L210W and D218E/D67N), and most of the time greater than 10. with the noticeable exception of I135L where only 2 known RAMs give ratios with lower CI bounds greater than 4. The ratios computed on the African dataset (S2 Fig) tell a similar story albeit with smaller ratio values due to a smaller number of occurrences of both new mutations and known RAMs.

The genetic barrier to resistance for each of these new mutations is quite low, with a minimum of 1 base change for each of them (Table 3). We also computed the average codon distance (i.e. number of different bases), weighted by the prevalence of wild and mutated codons at the given positions in the UK (Table 3) and Africa (Sup Mat) datasets, and in each case the average codon distance was always close to 1. In other words, at the DNA level these mutations are expected to be relatively frequent. However, their frequencies are much higher in treated/with-RAM sequences than in naive/without-RAM ones (Table 3). Moreover, if we look at the BLOSUM62 scores (Table 3) some of these mutations change radically the physicochemical properties of the amino acids, which reinforces again the likelihood that these mutations are associated to resistances.

To gain more insight on these new mutations we also observed their spatial location on the 3-D HIV-1 RT structure using PyMol [25]. HIV-1 RT is a heterodimer with two subunits translated from the same sequence with different lengths and 3-D structures. The smaller p51 subunit (440 AAs) has a mainly structural role, while the larger p66 (560 AAs) subunit has the active site at positions 110, 185 and 186. The p66 subunit also has a regulatory pocket behind the active site: the non-nucleoside inhibitor binding pocket (NNIBP) formed of several sites of the p66 subunit as well as site 138 of the p51 subunit. Nucleoside RT Inhibitors (NRTI) are nucleotide analogs and bind in the active site, blocking reverse transcription. Non-Nucleoside RT Inhibitors (NNRTI) bind in the NNIBP, changing the protein conformation and blocking reverse transcription. More details on the structure and function of HIV-1 RT can be found in [26]. A general view of where the new mutations are situated with regards to the other important sites of HIV-1 RT is shown in Fig. 4, and is detailled below.

**Fig 4.**
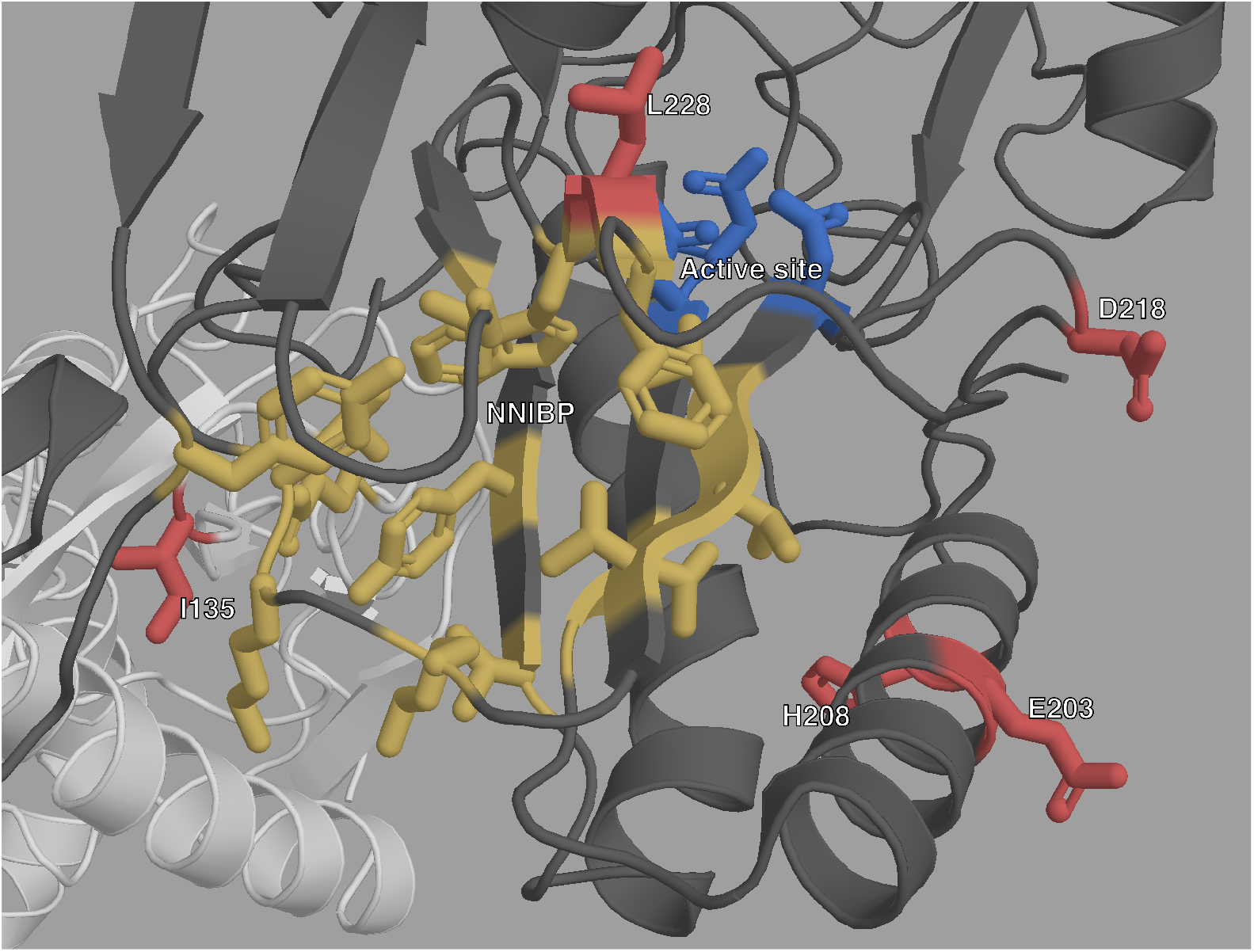
Structure of HIV-1 RT with highlighted important sites. The p66 subunit is colored dark gray and the p51 subunit white. The active site is highlighted in blue, and the NNIBP is highlighted in yellow. The sites of new mutations are colored in red.

### L228R / L228H

L228R is the most important of these new mutations according to the feature importance ranking done above. This is reflected in the very high over-representation in RTI-experienced sequences and sequences with known RAMs shown in Table 3. When looking at the detailed ratios shown in Fig. 3, we observe that L228R presents high ratio values with mainly NRTI RAMs, but also has high ratio values for NNRTI RAMs such as Y181C and L100I, and this is even more so for ratios computed on the African dataset (S2 Fig). L228H is very similar in all regards to L228R, however its highest ratios are exclusively with NRTI RAMs.

Site 228 of the p66 subunit is located very close to the active site of RT, where NRTIs operate (Fig. 4 and S4 Fig) which could explain the role that L228R and L228H seem to have in NRTI resistance. However, site 228 of the p66 subunit is also between sites 227 and 229 which are both part of the NNIBP. Furthermore, both L228H and L228R have very low BLOSUM62 score, of -3 and -2 respectively (Table 3). Arginine and Histidine (H) are both less hydrophobic that Leucine (L), and have positively charged side-chains. This important change in physicochemical properties could explain the role they both seem to have in NRTI resistance. However, while both Arginine and Histidine are larger than Leucine, Arginine is also fairly larger than Histidine, which is aromatic. This difference between both residues might indicate an association L228R seems to have with NNRTI resistance that L228H does not.

### E203K / H208Y

Both E203K and H208Y are highly over-represented in RTI-experienced sequences and sequences with known RAMs. They both have high ratio values for NRTI RAMs. Furthermore the most highly valued RAM ratios in Fig 3, are very similar for E203K and H208Y. Structurally they are close to each other on an alpha helix which is close to the active site.

Both E203K and H208Y have positive, albeit not maximal, BLOSUM62 scores, meaning they are fairly common substitutions. However, these mutations induce some change in physicochemical properties with Tyrosine (Y) being less polar than Histidine (H), and the change from Glutamic Acid (E) to Lysine (K) corresponding to a change from a negatively charged side chain to a positively charged one.

All this, combined with their structural proximity and the shared high ratio values for single RAMs, suggests a similar role in NRTI resistance.

### I135L

In Table 3 and Fig 3, we observe that I135L has the lowest ratio values of all the new mutations, with CI bounds lower than 2 in Table 3’s general ratios. However, it is the most prevalent of the new mutations. If we look at the detailed ratios of Fig 3, we see that I135L is significantly over-represented in sequences with NNRTI RAMs, specifically A98G and P225H. Structurally this makes sense: On the p66 subunit, site 135 is on the outside, far from both the active site and the NNIBP. However, site 135 on the p51 subunit is located very close to the NNIBP (Fig 3, and S3 Fig).

The BLOSUM62 score for this substitution is quite high (Table 3), which is expected since both residues are very similar to one another, differing only by the positioning of one methyl group. However, Leucine (L) is less hydrophobic than Isoleucine (I), despite they are still both classified as hydrophobic residues (S5 Table).

The proximity between site 135 and the pocket in which NNRTI RAMs bind, as well as the high ratio values for these NNRTI RAMs leads us to believe that I135L could play a subtle accessory role in NNRTI resistance, either by enhancing the effect of some NNRTI RAMs (typically, A98G and P225H), or by compensating for loss of fitness.

### D218E

D218E is also highly over-represented in both RTI-experienced sequences and sequences with known RAMs. It has infinite ratio values in the African dataset (Table 3), because it is quite rare in this dataset, and all of its 25 occurrences are in sequences that have at least one known RAM and are RTI-experienced. In fact, from the UK dataset we can see that D218E has some of the highest ratio values for individual RAMs (along with H208Y). The majority of these very high ratio values occur for NRTI RAMs. Site 218 on the p66 subunit is quite close to the RT active site, which could explain the role D218E seems to have in NRTI resistance. Aspartic acid (D) and Glutamic acid (E) are very similar amino acids, both acidic with negatively charged side-chains, as reflected in their fairly high BLOSUM62 score, the main difference between both being molecular weight, with E being slightly larger than D.

## Discussion and perspectives

Our method has allowed us to identify six mutations that might play a role in drug resistance in HIV. These mutations are significantly over-represented in RTI-experienced sequences, as well as sequences exhibiting at least one other known RAM. The fact that models trained on the UK are still performant on such a different dataset as the African one proves that the learned classifier models have acquired generalized knowledge on resistance. For all of these new mutations their spatial positioning on HIV-1 RT is consistent with our conclusions, as all were either close to the active site or the regulatory binding pocket.

Some of the mutations we have identified as potentially associated with resistance have been mentioned in previous studies. L228R/H have been observed before [27] and were suggested to be associated with reduced susceptibility to didanosine [28, 29]. I135L has been observed in sequences with reduced susceptibility to NNRTIs [30]. H208Y has been associated with NNRTI and NRTI resistance [31] and it has been suggested that it has an accessory role in NRTI resistance [32]. E203K, D218E, L228RH and H208Y have all been mentioned in [33] as probably linked to phenotypic resistance to NRTI and NNRTI.

However, none of these mutations has been experimentally confirmed as conferring or helping with drug resistance to the best of our knowledge. The fact that we find them again with a big data analysis of highly different sequences and involved statistical selection procedure combining multiple testing and machine learning, and that we have very high significance clearly indicates their potential role in resistance. Therefore, we believe they are sufficiently linked to drug resistance that they garner a closer inspection either in-vitro or in-vivo to determine the mechanisms that could allow them to play a role in resistance.

By using machine learning classifiers that can detect interactions between features we should be able to detect epistasis as we defined it above. However, by treating the Fisher exact tests as the internal model for a very simple baseline classifier, and observing little performance gain for more sophisticated classifiers, it is likely that epistasis plays a limited role in drug resistance in HIV-1 RT, beyond the standard scheme where DRMs are associated to auxiliary and compensatory mutations. However, one advantage of machine learning classifiers, is that they are probabilistic, meaning that they can give more nuanced insights into the nature or resistance level of a given sequence than the classical binary presence/absence of RAMs approach. In this regard logistic regression appears as a method of choice, showing similar or better performance than other classifiers, and an easy interpretation that is facilitated by the lasso regularization which performs a simple feature selection and retains the most important ones. Similar results were already observed on other sequence analysis tasks [34].

Even though we did not find any evidence of epistasis, with the now used pluripotent therapeutic strategies, there might be some epistatic effects with other regions of the HIV genome that are targeted by some of the drugs. These potential effects however, lie outside the scope of this study.

Because of the lack of detailed treatment history metadata, we did not distinguish mutations arising from NRTIs or NNRTIs. We believe that a large amount of high quality sequence data, along with a sufficiently detailed log of treatments and drugs the sequences were exposed to, could allow us to use our machine-learning approach to find mutations related to specific drugs and thus furthering our knowledge of HIV drug resistance, giving clinicians more tools to manage and help infected patients.

## Acknowledgments

We thank Anna Zhukova, Frédéric Lemoine and Marie Morel for their help and suggestions.

## Funding

This work was supported by the EU-H2020 Virogenesis project (grant number 634650, to L.B. and O.G.) and PRAIRIE (ANR-19-P3IA-0001).

We also thank the UK HIV Drug Resistance Database and the UK Collaborative HIV Cohort:

## Steering committee

David Asboe, Anton Pozniak (Chelsea & Westminster Hospital, London); Patricia Cane (Public Health England, Porton Down); David Chadwick (South Tees Hospitals NHS Trust, Middlesbrough); Duncan Churchill (Brighton and Sussex University Hospitals NHS Trust); Simon Collins (HIV i-Base, London); Valerie Delpech (National Infection Service, Public Health England); Samuel Douthwaite (Guy’s and St. Thomas’ NHS Foundation Trust, London); David Dunn, Kholoud Porter, Anna Tostevin, Oliver Stirrup (Institute for Global Health, UCL); Christophe Fraser (University of Oxford); Anna Maria Geretti (Institute of Infection and Global Health, University of Liverpool); Rory Gunson (Gartnavel General Hospital, Glasgow); Antony Hale (Leeds Teaching Hospitals NHS Trust); Stéphane Hué (London School of Hygiene and Tropical Medicine); Michael Kidd (Public Health England, Birmingham Heartlands Hospital); Linda Lazarus (Expert Advisory Group on AIDS Secretariat, Public Health England); Andrew Leigh-Brown (University of Edinburgh); Tamyo Mbisa (National Infection Service, Public Health England); Nicola Mackie (Imperial NHS Trust, London); Chloe Orkin (Barts Health NHS Trust, London); Eleni Nastouli, Deenan Pillay, Andrew Phillips, Caroline Sabin (University College London, London); Kate Templeton (Royal Infirmary of Edinburgh); Peter Tilston (Manchester Royal Infirmary); Erik Volz (Imperial College London, London); Ian Williams (Mortimer Market Centre, London); Hongyi Zhang (Addenbrooke’s Hospital, Cambridge).

## Coordinating Center

Institute for Global Health, UCL (David Dunn, Keith Fairbrother, Anna Tostevin, Oliver Stirrup)

## Centers contributing data

Clinical Microbiology and Public Health Laboratory, Addenbrooke’s Hospital, Cambridge (Justine Dawkins); Guy’s and St Thomas’ NHS Foundation Trust, London (Emma Cunningham, Jane Mullen); PHE – Public Health Laboratory, Birmingham Heartlands Hospital, Birmingham (Michael Kidd); Antiviral Unit, National Infection Service, Public Health England, London (Tamyo Mbisa); Imperial College Health NHS Trust, London (Alison Cox); King’s College Hospital, London (Richard Tandy); Medical Microbiology Laboratory, Leeds Teaching Hospitals NHS Trust (Tracy Fawcett); Specialist Virology Centre, Liverpool (Elaine O’Toole); Department of Clinical Virology, Manchester Royal Infirmary, Manchester (Peter Tilston); Department of Virology, Royal Free Hospital, London (Clare Booth, Ana Garcia-Diaz); Edinburgh Specialist Virology Centre, Royal Infirmary of Edinburgh (Lynne Renwick); Department of Infection & Tropical Medicine, Royal Victoria Infirmary, Newcastle (Matthias L Schmid, Brendan Payne); South Tees Hospitals NHS Trust, Middlesbrough (David Chadwick); Department of Virology, Barts Health NHS Trust, London (Mark Hopkins); Molecular Diagnostic Unit, Imperial College, London (Simon Dustan); University College London Hospitals (Stuart Kirk); West of Scotland Specialist Virology Laboratory, Gartnavel, Glasgow (Rory Gunson, Amanda Bradley-Stewart).

The UK HIV Drug Resistance Database and the UK Collaborative HIV Cohort are currently supported by the UK Medical Research Council (Award Number 164587).

## Supporting information

**S6 File. List of known DRMs**. A .csv file containing all the known RAMs used in this project as well as the corresponding feature name in the encoded datasets. Obtained from (hivdb.stanford.edu/dr-summary/comments/NRTI/) and (hivdb.stanford.edu/dr-summary/comments/NNRTI/).

**S7 Web-page. Code repository for this project**. The different classes and methods used in the study’s pipeline were organized in a python module that can be accessed at github.com/lucblassel/utils hiv. The pipeline as well as additional tab delimited files can be found at github.com/lucblassel/HIV-DRM-machine-learning.

## S1 Appendix Technical appendix

### Data

#### Data Availability

The policy of the UK HIV Drug Resistance Database is to make information available to any bona fide researcher who submits a scientifically robust proposal, provided data exchange complies with Information Governance and Data Security Policies in all the relevant countries. This includes replication of findings from published studies, although the researcher would be encouraged to work with the main author of the published paper to understand the nuances of the data. Enquiries should be addressed to in the first instance. More information on the UK dataset is also available on the UK CHIC homepage: www.ukchic.org.uk

The West and central African dataset is available as supplementary information along with a metadata file containing HIV subtype, treatment information and known RAM presence/absence for each sequence. A similar metadata file, containing the same information is also made available for the UK dataset.

Predictions made for each sequence of both datasets, by all of the trained classifiers are made available on the folloqing GitHub repository: <u>github.com/lucblassel/HIV-DRM-machine-learning</u>, as well as the importance values for each mutation and each trained classifier.

#### Data Preprocessing

For both the African and UK datasets, the sequences were truncated to keep sites 41 to 235 of the RT protein sequence before encoding. This truncation was needed to avoid the perturbation to classifier training due to long gappy regions at the beginning and end of the UK RT alignment caused by shorter sequences. These positions were determined with the Gblocks software [35] with default parameters, except for the Maximum number of sequences for a flanking position, set to 50,000, and the Allowed gap positions, which was set to “All”. The encoding was done with the OneHotEncoder from the category-encoders python module [36].

#### Classifiers

We used classifier implementations from the scikit-learn python library [37], `RandomForestClassifier` for the random forest classifier, `MultinomialNB` for Naïve Bayes and `LogisticRegressionCV` for logistic regression.

`RandomForestClassifier` was used with default parameters except:

- `“n jobs”=4`
- `“n estimators”=5000`

`LogisticRegressionCV` was used with the following parameters:

- `“n jobs”=4`
- `“cv”=10`
- `“Cs”=100`
- `“penalty”=‘l1’`
- `“multi class”=‘multinomial’`
- `“solver”=‘saga’`
- `“scoring”=‘balanced accuracy’`

`MultinomialNB` was used with default parameters.

For the Fisher exact tests, we used the implementation from the scipy python library [38], and corrected p-values for multiple testing with the statsmodels python library [39] using the `“Bonferroni”` method.

#### Scoring

To evaluate classifier performance several measures were used. We computed balanced accuracy instead of classical accuracy, because it can be overly optimistic, especially when assessing a highly biased classifier on an unbalanced test set [23].The balanced accuracy is computed using the following formula, where *TP* and *TN* are the number of true positives and true negatives respectively, and *FP* and *FN* are the number of false positives and false negatives respectively:

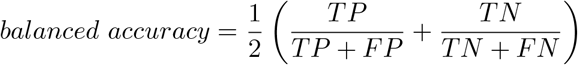

We also computed adjusted mutual information (AMI). We chose it over mutual information (MI) because it has an upper bound of 1 for a perfect classifier and is not dependent on the size of the test set, allowing us to compare the performance for differently sized test sets [24]. The adjusted mutual information of variables *U* and *V* is defined by the following formula, where *MI*(*U, V*) is the mutual information between variables *U* and *V, H*(*X*) is the entropy of the variable *X* (= *U* or *V*) and *E{MI*(*U, V*)*}* is the expected MI, as explained in [40].

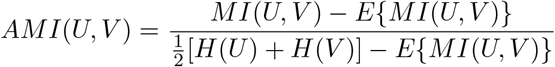

MI was used to compute the *G* statistic, which follows the chi-square distribution under the null hypothesis [41]. This was used to compute p-values for each of our classifiers and assess the significance of their performance. *G* is defined by equation below, where *N* is the number of samples.

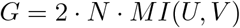

Finally, to check the probabilistic predictive power of the classifiers we also computed the Brier score which is the mean squared difference between the ground truth and the predicted probability of being of the positive class for every sequence in the test set (therefore lower is better for this metric). The Brier score is defined in equation below, where *p*_*t*_ is the predicted probability of being of the positive class for sample *t* and *o*_*t*_ is the actual class (0 or 1, 1=positive class) of sample *t*:

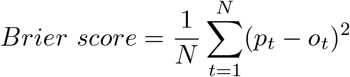

We used the following implementations from the scikit-learn python library [37] with default options:

- `balanced accuracy score`
- `mutual info score`
- `adjusted mutual info score`
- `brier score loss`

We used the following ratio to observe the relationship between one of our new mutations and a binary character *X* such as treatment status or presence/absence of a known RAM.

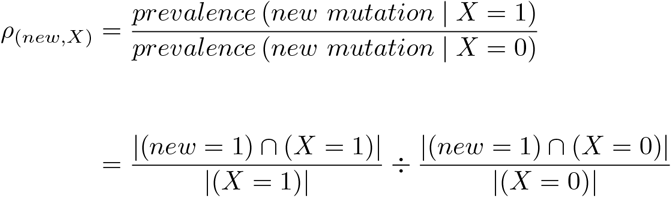

---

**Fig S2.**
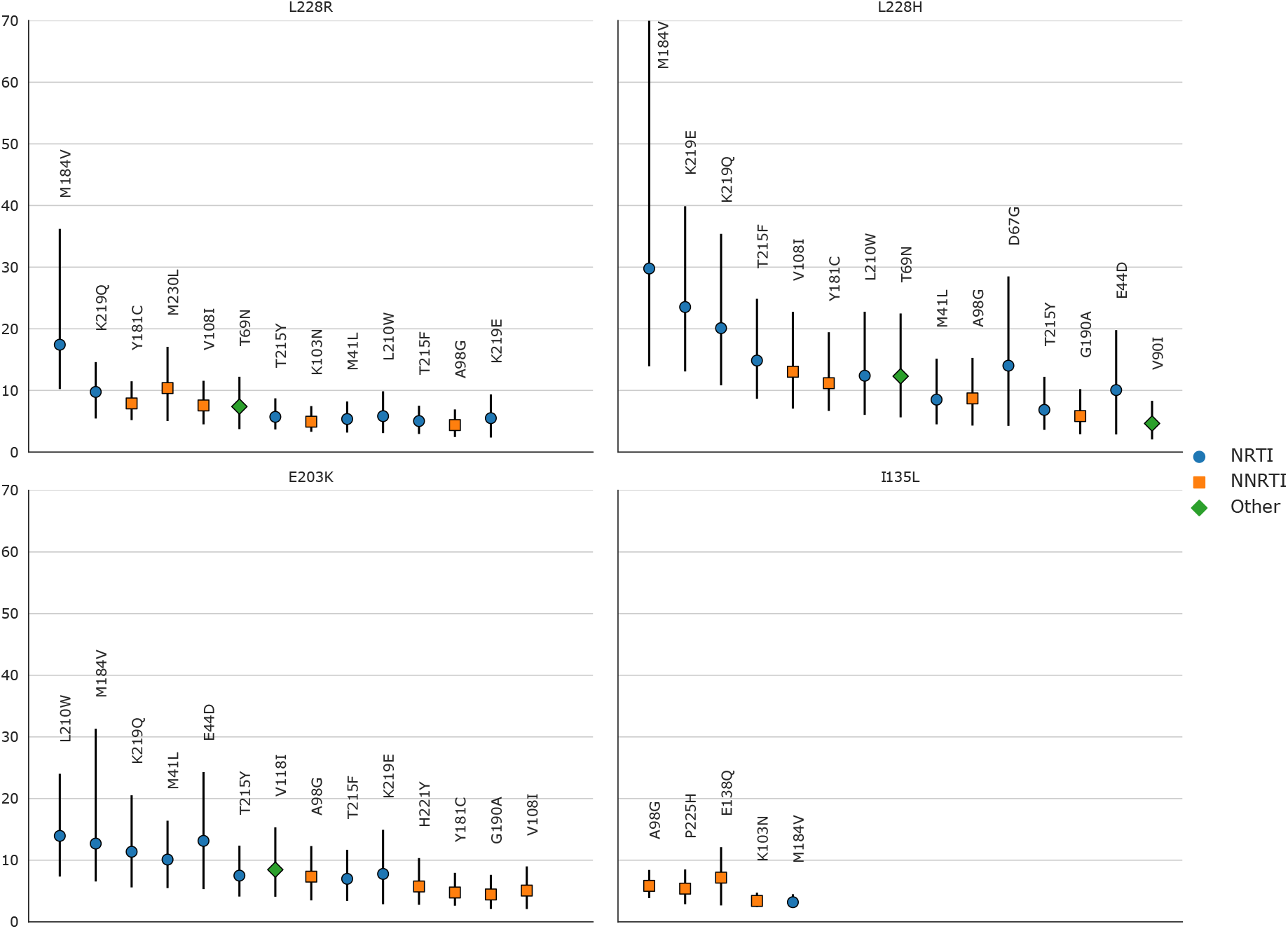
Prevalence ratios of the new mutations with regards to known RAMs on the African dataset. (i.e. the prevalence of the new mutation in sequences with a given RAM divided by the prevalence of the new mutation in sequences without the RAM). Ratios were only computed for mutations (new and RAMs) that appeared in at least 30 sequences, which is why ratios were not computed for H208Y and D218E. 95% confidence intervals, represented by vertical bars, were computed with 1000 bootstrap samples of the African sequences. Only ratios with a lower CI boundary greater than 2 are shown. The shape and color of the point represents the type of RAM as defined by Stanford’s HIVDB. Blue circle: NRTI, orange square: NNRTI, green diamond: Other. For the prevalence ratio of L228H with regards to M184V, the upper CI bound is infinite. The new RAMs have high ratio values for known RAMs similar to those obtained on the UK dataset. We also arrive at similar conclusions, I135L being associated with NNRTIs, E203K and L228H to NRTI and L228R to both. Ratio values are shown from left to right, by order of decreasing values on the lower bound of the 95% CI.

**Fig S3.**
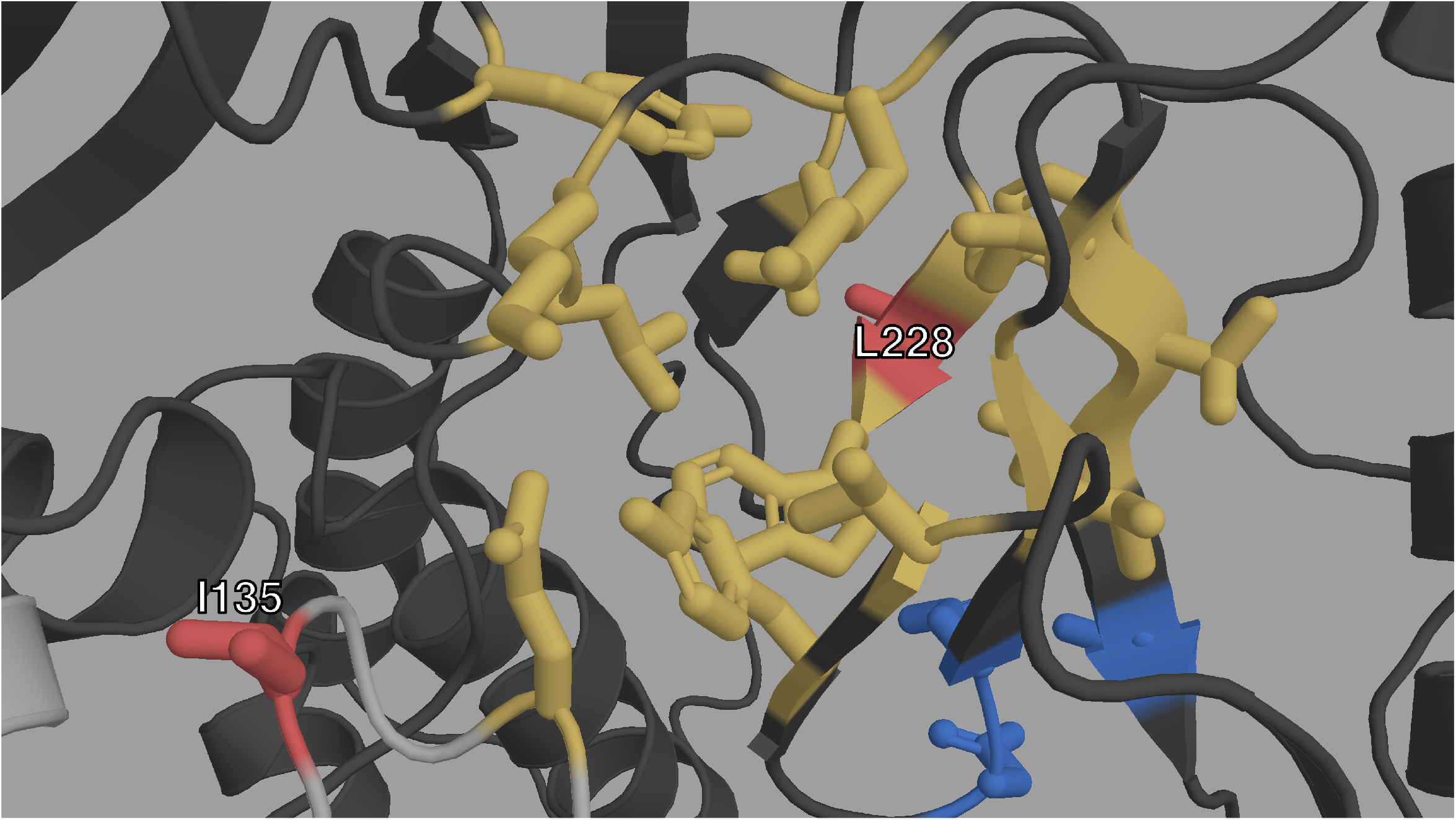
Closeup structural view of the entrance of the NNIBP of HIV-1 RT. The p66 subunit is colored in dark gray, the p51 subunit in light gray. The NNIBP is highlighted in yellow. The active site is colored in blue. We can see the physical proximity of I135 (red) to the entrance of the NNIBP. We can also see how L228 (red) is between 2 AAs of the NNIBP.

**Fig S4.**
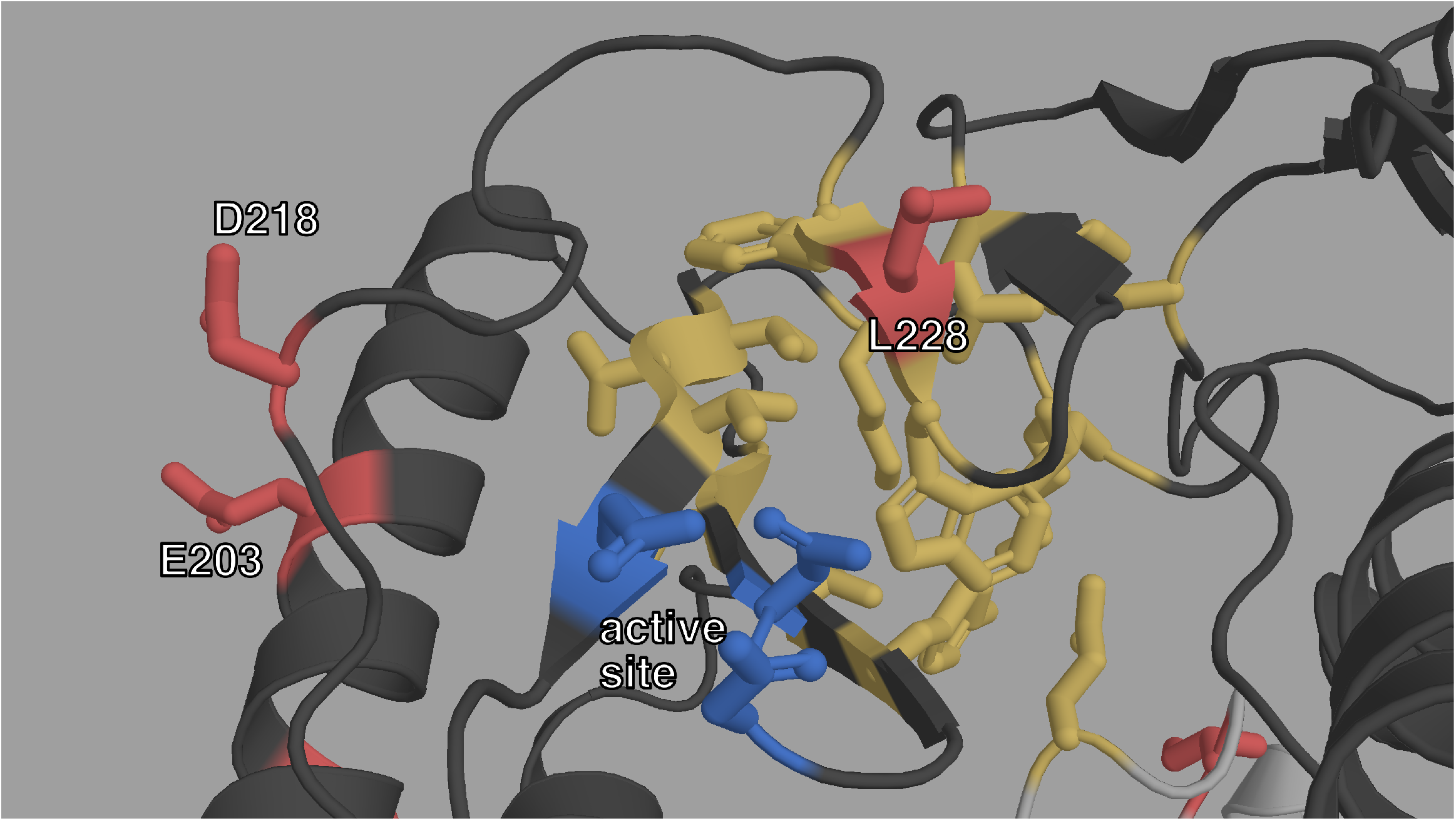
Closeup structural view of the active site of HIV-1 RT. The p66 subunit is colored in dark gray, the p51 subunit in light gray. The active site is highlighted in blue. The NNIBP is colored in yellow. L228, E203 and D218 (red) are also very close on either side of the active site.

**Table S5.**
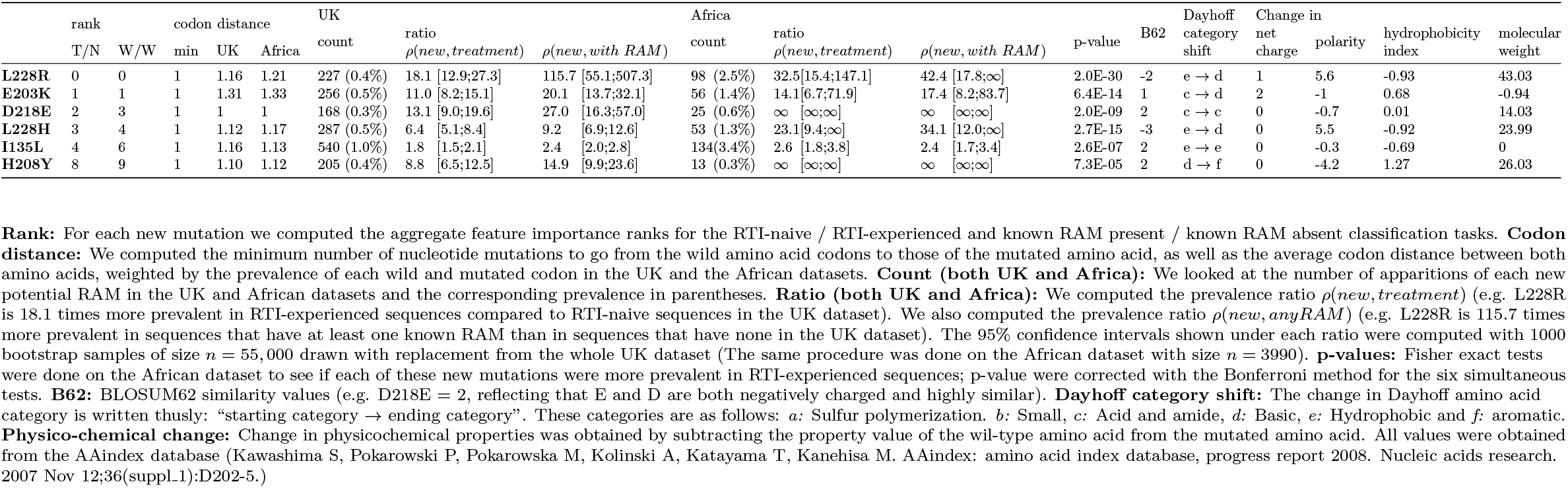
Detailed view of the characteristics of new potential RAMs

